# Cyclic nature of the REM-like state in the cuttlefish *Sepia officinalis*

**DOI:** 10.1101/224220

**Authors:** T. L. Iglesias, J. G. Boal, M.G. Frank, J. Zeil, R.T. Hanlon

## Abstract

Sleep is a state of immobility characterized by three key criteria: an increased threshold of arousal, rapid reversal to an alert state, and evidence of homeostatic “rebound sleep” in which there is an increase of time spent in this quiescent state following sleep deprivation. Common European cuttlefish, *Sepia officinalis*, show states of quiescence during which they meet the last two of these three criteria, yet also show spontaneous bursts of arm and eye movements that accompany rapid changes in chromatophore patterns in the skin. Here we report that this rapid-eye-movement (REM)-like state is cyclic in nature. Iterations of the REM-like state last 2.42 ± 0.22 min (±SE) and alternate with 34.01 ± 1.49 min of the quiescent sleep-like state. These states alternate for durations lasting 176.89 ± 36.71 min. We found clear evidence that this REM-like state (i) occurs in animals younger than previously reported; (ii) follows an ultradian pattern; (iii) includes intermittent dynamic chromatophore patterning, representing fragments of normal patterning seen in the waking state for a wide range of signaling and camouflage; and (iv) shows variability in the intensity of expression of these skin patterns between and within individuals. These data suggest that cephalopods, which are molluscs with an elaborate brain and complex behavior, possess a sleep-like state that resembles behaviorally the vertebrate REM sleep state, although the exact nature and mechanism of this form of sleep may differ from that of vertebrates.

## Introduction

In mammals, birds, and some reptiles (Shein-Idelson, Ondracek, Liaw, Reiter, & Laurent, 2016) (but see (Aulsebrook, Jones, Rattenborg, Roth, & Lesku, 2016)) sleep is characterized neurologically by the occurrence of two distinct brain states denoting slow-wave sleep and rapid-eye-movement (REM) sleep. Behaviorally, slow-wave sleep is quiescent, while REM sleep is accompanied by rapid eye movements and muscular atonia with occasional muscular twitches (Mascetti, 2016; Stickgold & Walker, 2010). Sleep follows a circadian rhythm and within each sleep bout, slow-wave sleep and REM sleep follow in an ultradian rhythm (Beckers & Rattenborg, 2015; Low, Shank, Sejnowski, & Margoliash, 2008; Martinez-Gonzalez, Lesku, & Rattenborg, 2008; Shein-Idelson et al., 2016; I. Tobler, 1995). Slow-wave sleep alternates with REM sleep and multiple iterations of REM sleep can occur within one bout of sleep, with slow-wave sleep always preceding REM sleep (Stickgold & Walker, 2010). The duration of different sleep states and frequency at which these states alternate *(i.e*. the periodicity), varies across species (John A. Lesku, Roth, Amlaner, & Lima, 2006). Periodicity is widely variable in mammals; for example, humans have a 90-minute periodicity whereas rats have a ten-minute periodicity (Stickgold & Walker, 2010; I. Tobler, 1995). Birds have short sleep periodicities that may last less than a minute, with iterations of REM sleep just 2-10 seconds in duration (J. A. Lesku & Rattenborg, 2014; Stickgold & Walker, 2010), or up to five minutes in ostriches (John A. Lesku et al., 2011), and slow-wave sleep intervals that range from 10-100 seconds (Low et al., 2008; Martinez-Gonzalez et al., 2008). The recent discovery of REM sleep in a reptile revealed an 80-second sleep periodicity (Shein-Idelson et al., 2016). An ultradian rhythm, therefore, appears to be an inherent characteristic of sleep in organisms that experience multiple stages of sleep (Beckers & Rattenborg, 2015; Low et al., 2008; Martinez-Gonzalez et al., 2008; I. Tobler, 1995).

The cuttlefish, *Sepia officinalis* (Figure 1), demonstrates several behavioral indicators that its quiescent sleep-like (QS) state may be analogous to sleep in other organisms (Frank, Waldrop, Dumoulin, Aton, & Boal, 2012). For example, cuttlefish show behaviors that accompany a state of immobility, such as rapid reversal to an alert state, and QS-deprivation induces a homeostatic response known as “rebound sleep” (Frank et al., 2012). The third criterion, an increase in arousal threshold during the sleep state, has not yet been successfully examined in this species; however, all three characteristics have been observed in another cephalopod, *Octopus vulgaris* (Brown, Piscopo, De Stefano, & Giuditta, 2006; Meisel, Byrne, Mather, & Kuba, 2011) and in many other invertebrates (for example: fruitflies, bees, jellyfish, worms, and crayfish respectively; Cirelli & Bushey, 2008; Kaiser, 1988; Nath et al., 2017; Raizen et al., 2008; Ramon, Hernandez-Falcon, Nguyen, & Bullock, 2004). In this study, we video recorded adult, non-senescent, cuttlefish day and night to further examine the REM-like behavior reported by Frank et al. (2012) in adult, senescing cuttlefish. In the present study, multiple iterations of the REM-like state were observed in adult, non-senescing animals that were in the QS state and the behaviors were as described in Frank et al. (2012): characterized by rapid chromatophore changes (skin brightness and patterning), skin-texture changes, rapid eye-movement, and arm twitching. Our aim here was to determine if there was evidence of an ultradian rhythm — typical periodic cycle between slow-wave sleep and REM — using the observable state of rapid chromatophore changes as a proxy for the REM-like state. Additionally, using measures of pixel intensity, we compared chromatophore changes within and between individuals to assess if chromatophore patterns followed a stereotyped sequence of activation within and/or between individuals.

**Figure 1.**
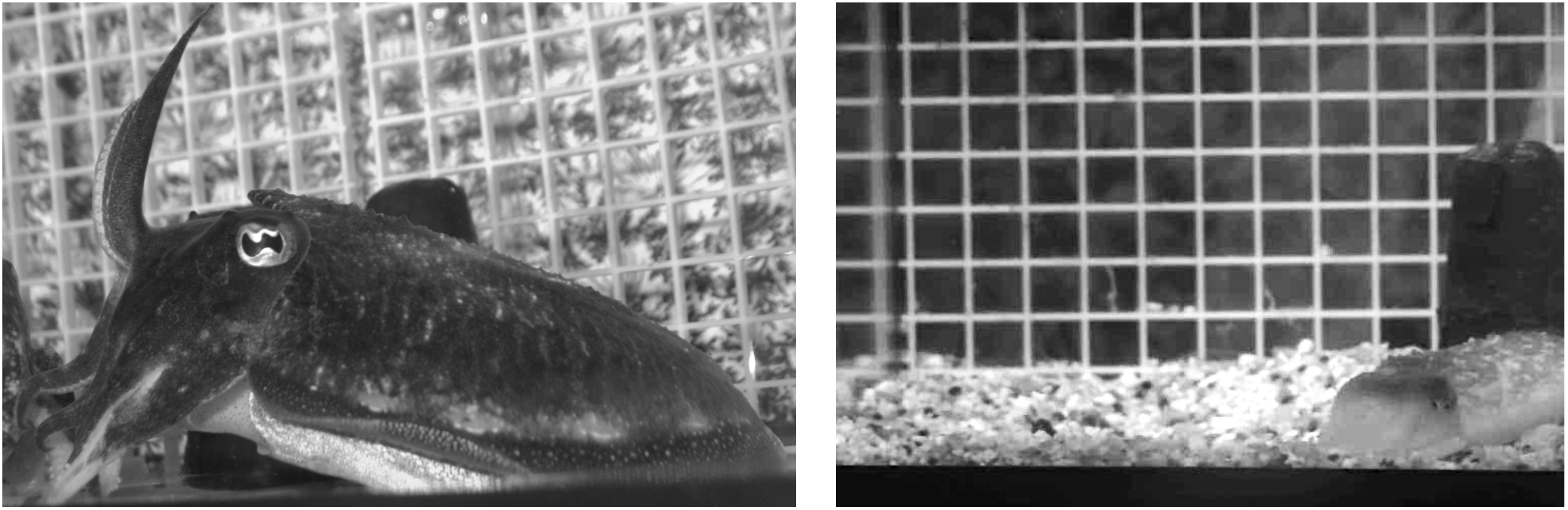
Cuttlefish in awake state with pupils dilated (left). Cuttlefish in quiescent sleep-like (QS) state, partly buried in substrate with pupils closed (right).

## Methods

Six common cuttlefish, one-year-old *Sepia officinalis* (ca. 7.5cm mantle length), hatched from eggs collected from the English Channel and raised in captivity in the Marine Resources Center at the Marine Biological Laboratory, were transferred singly to one of four 19-liter glass tanks (L x W x D: 40.64cm, 20.32cm, 24.5cm). Visual isolation was accomplished by covering the sides of the tanks with black and white patterned cloth, and white grating was used to constrain cuttlefish to one part of the tank for better video viewing (see Figure S1). Animals had ample room to swim (L x W x D: 24.5cm, 20.32cm, 24.5cm). Black plastic sheeting was used to isolate the tank area and reduce human disturbance. At least 1cm of gravel substrate was provided as this seemed to calm the animals, along with a 4cm tall, black cone placed in front of the inflow tube to reduce water agitation and cuttlefish movement due to water flow (Figure S1). Animals were provided thawed frozen shrimp every day. Tanks were supplied with flow-through seawater at a temperature of 15°C. At this temperature, animals grew slowly and did not develop secondary sex characteristics; thereby precluding sex-determination of animals used in this study. Room lighting was on a 12-hour light: 12-hour dark time cycle. One overhead camera and one side-view camera, Sanyo CCD camera model VCB-3384 and Everfocus Polestar II model EQ610, were placed such that they could record activity in one tank at a rate of 30 frames per second in black and white. Red lights (to which cuttlefish exhibited rapid pupil expansion indicating weak sensitivity) were in constant use to illuminate the dark period to enable video recording of body patterns from above, and eye movement and arm twitching from the side. Video was captured and saved in real time to a hard drive.

Video was reviewed at high speed (4X-32X) approximately every 24-48 hours to determine if an animal had experienced a REM-like state. After REM-like behavior was recorded at least once, the animal was returned to the regular holding tanks and a new individual was placed in that observation tank. The camera pair was moved to record an animal that had already spent at least 24 hours in an observation tank. The number of days of video that were collected for each animal was as follows: Hippo (28), zZ (12), MrChips (9), Medusa (9), Dumbo (7), and Clyde (5).

Durations of REM-like and QS states were determined from the video using the video time code for each recording. Video clips were made of the REM-like states including a 30 s lead-in of QS state before the start of the REM-like state and ending approximately 5 s after the animal expelled water through the siphon and/or repositioned itself on the substrate (Figure 2). These behaviors were performed consistently after the REM-like state. Following these movements, animals either resumed the QS state or became active with open pupils. The pixel intensity data for this 30 s lead in were excluded when calculating the distance matrix for the hierarchical clustering dendrogram and for the Mantel tests (see below). Overhead and side-view camera footage of REM-like states were combined in one video using Adobe After Effects (Figure 2, Video S1-S5) so that QS states could be confirmed (animal partially buried in substrate with pupils closed (Frank et al., 2012)).

**Figure 2.**
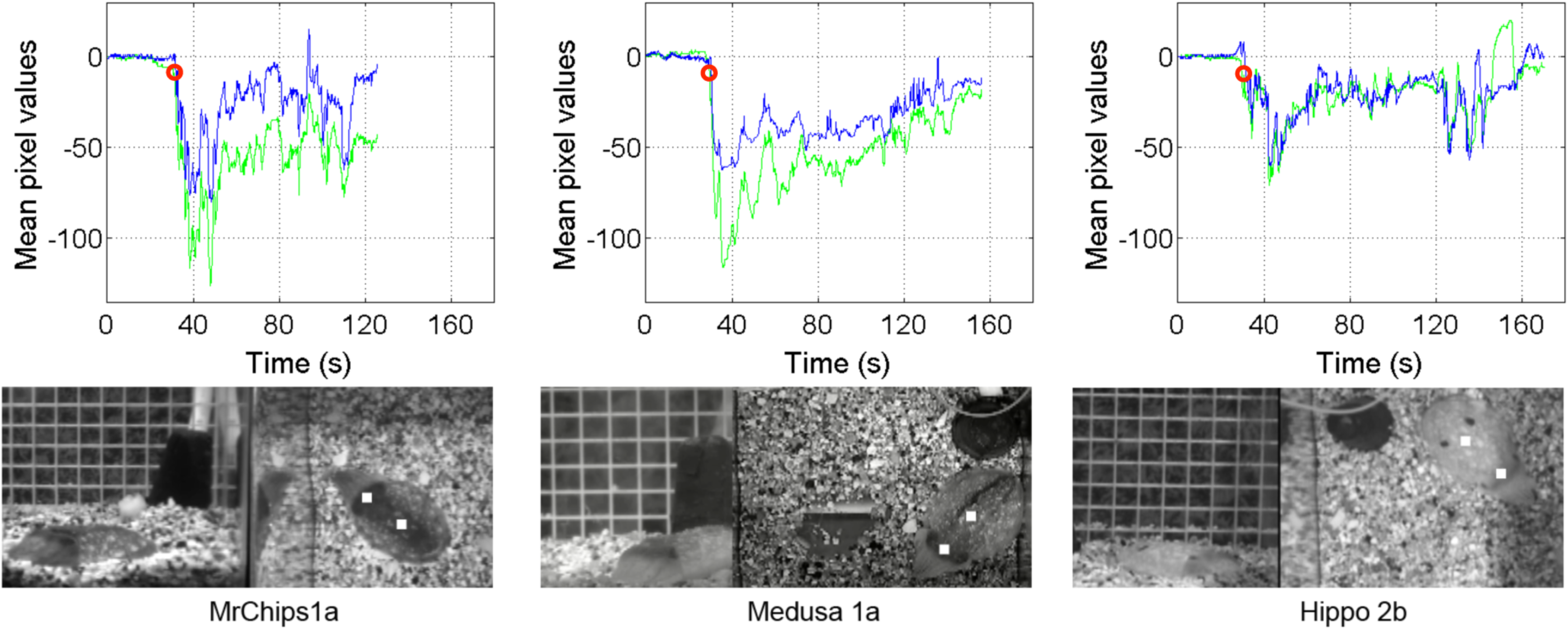
Stills of video analyzed next to pixel intensity measures for three cuttlefish (MrChips, Medusa, and Hippo). The white squares indicate regions of interest measured. The graphs show the pixel intensity measures for head and mantle regions for these video clips (head is the green line, mantle is blue line). The first 30 seconds of pixel intensity at the beginning is the QS state where pixel intensity was standardized at 0 to allow comparisons. Lower pixel values (more negative) indicate darker pigmentation. Our criterion for the start of a REM-like state iteration is marked by a red circle (rapid darkening of pigmentation in both regions by 10 mean pixel values).

Pixel value measurements were used to assess chromatophore activity; they were performed using custom-written MATLAB code (MathWorks, Natick, MA, USA). Most animals were relatively immobile as they were slightly buried in the substrate; however, one animal, zZ, drifted during the REM-like state because the substrate was too shallow as a result of repeated self-burying in the same spot. Videos for zZ were digitized using Adobe After Effects, using the motion-tracking feature so that the region of interest (head or mantle) was constantly at the center of view. Video clips of REM-like states were analyzed in MATLAB. A square 5 x 5 pixel region was selected on the head and on the mantle of the cuttlefish (Figure 2). The average pixel values (ranging from 0 to 255 in 8bit grey level images) were collected for each video frame (30fps) over the entire video clip for these two regions (Figure 2).

The absolute pixel values were affected by the amount of ambient light in the video. Because the aim was to compare patterns of pixel value changes throughout the REM-like state within and between individuals and across different iterations of the REM-like state, the pixel value at the beginning of sequences was subtracted from all subsequent values so that all REM-like state iterations had a standardized start at zero (see Figure 2). In other words, in the standardized data, positive numbers indicate brightening of pixels and negative numbers indicate darkening of pixels. The REM-like state began with a rapid darkening of the head and mantle region. We visually determined from the videos that a darkening by approximately 10 mean pixel values coincided with the start of the REM-like state and only used the subsequent sequence for our analysis (see Figure 2).

The temporal structure of body patterning was compared between and within individuals using the R package *dtwclust* (Sarda-Espinosa, 2016) which calculated all pair-wise distances/dissimilarities between pixel value series for all 55 iterations of the REM-like state. This package stretches or condenses signals, based on user specified settings, to determine how similar (or different) signals are to one another (Figure S2). The settings disallowing unlimited stretching and condensing of signals were determined using the R package *dtw* (Giorgino, 2009) and were as follows: step=symmetric2, window.type=slantedband, and window.size=30. In *dtwclust* the function ‘hclust’ was used to build a hierarchical dendrogram for head and mantle data with animal and iteration of REM-like state annotated at the tips to visually assess clustering. The R package *ape* (Paradis, Claude, & Strimmer, 2004) was used to format the hierarchical dendrogram.

The R package *Ecodist* (Goslee & Urban, 2007) was used to perform Mantel tests between the pixel intensity distance matrix and binary matrices for animal identity and REM-like state iteration. Each binary matrix was compared against the pixel distance matrix independently. All Mantel tests were performed using 100,000 permutations, sampling with replacement, and 100,000 iterations for bootstrapped confidence intervals. A Mantel correlogram was examined to understand the structure of the relationship when the Mantel r statistic was significant.

The duration of the REM-like state varied between individuals and REM-like state iteration, resulting in pixel intensity series of differing lengths; however, interpolating data onto the ends of the series to make them of equal length did not change results. Interpolated data were not used, therefore.

## Results

In total, 55 iterations of the REM-like state were observed. All six animals were recorded on camera demonstrating at least one sleep-like bout with the REM-like state alternating with the QS state. The six animals, with the numbers of sleep-like bouts and REM-like state iterations for each animal, were Hippo (2,15), zZ (1,10), MrChips (2,6), Medusa (3,11), Dumbo (1,9), and Clyde (1,4). Duration of the REM-like state iteration was on average 2.42 ± 0.22 min (±SE) and the time between REM-like state iterations *(i.e*. QS state intervals) was 34.01 ± 1.49 min (Table S1). We defined the periodicity as the time between onset of one REM-like state iteration to the onset of the next REM-like state iteration (marked by eye squinting, discussed below); this resulted in a 36.34 ± 1.46 min periodicity. On average, animals spent 13.36 min total (range = 1.93 - 23.07 min) in the REM-like state during a 176.89 ± 36.71 min sleep bout.

We observed eye and arm movements by every individual and in 54 out of 55 iterations (observation of the 55^th^ iteration was obscured by condensation on the tank). Eye movements and concurrent chromatophore patterning around the eyes and brain (collectively known as the “head” in cephalopods) included peculiar features that best characterize the REM-like state of *Sepia officinalis*. The eyes shifted fore and aft (or left and right when viewed from the side) in all animals, and the W-shaped slit pupil sometimes dilated to a round shape. Yet the more common eye behavior was a gross “squinting” of the whole eye socket in which various eye muscles contracted and squeezed the surrounding tissue to occlude the pupil. A close-up example of these strong eye movements can be seen in Video S1. Often this behavior was accompanied by sudden darkening of all the chromatophores around each eye and between the eyes, forming a “dark head bar” that has not been seen in normally behaving cuttlefish during daytime (Hanlon & Messenger, 1988). Sometimes the whole head was flattened as if the cuttlefish was ducking. Collectively, these represent conspicuous dramatic behaviors that are rarely or never seen in day-active cuttlefish; that is, they are REM-like state specific and their order of appearance was not fixed but variable. Animals were all observed to show a white head bar just prior to the REM-like state and perform this gross squinting, which coincided with the rapid chromatophore darkening at the start of the REM-like state. The eye squint was used to mark the start of the REM-like state for video editing and duration measurements.

Arm movements included slight twitching, raising of the 1^st^ pair, and movement of all arms (see Videos S4, S5 for full arm movement and arm raising). The only arm movement observed in all 55 iterations was slight arm twitching.

The videos revealed episodic expression of several of the 34 chromatic components of body patterns that *Sepia officinalis* is known to use for signaling and camouflage (Video S6) (Hanlon & Messenger, 1988). For example, cuttlefish showed each of the following: (i) mottle patterns (Chiao et al., 2009), a primary defense (camouflage); (ii) various expressions of deimatic (threat, startle) patterns, such as the uni-or bilateral paired mantle spots, which are secondary defenses; (iii) and dark flashing of the mantle or whole body, or the Passing Cloud pattern, which are examples of protean defense. These expressions were transitory and were not usually a full or normal expression of the patterns as seen in field or laboratory behavioral studies (Adamo, Kelly, Cheryl, & Ivy, 2006; Hanlon & Messenger, 1988, 2017; Langridge, 2006; Langridge, Broom, & Osorio, 2007; Staudinger et al., 2013).

To quantify the variability of skin chromatophore patterning activity (without tabulating species-specific displays), the distance matrix containing all pair-wise comparisons of REM-like state iterations was visualized in a hierarchical clustering dendrogram with animal ID and iteration tip labels for the two body regions (head - Figure 4; and mantle - Figure S3). No robust clustering by animal or by iteration (1^st^, 2^nd^. 3^rd^… REM-like state iteration within a cycle) was found for either body region. A Mantel statistical test comparing the pixel series distance matrix to a binary matrix for the head region showed a weak effect of animal identity: Mantel r = 0.09, two-tailed p < 0.05, and 95% confidence interval for Mantel r of [0.06, 0.14]. Results for pixel intensity distance data from the mantle of the cuttlefish were similar but not significant with Mantel r = 0.07, p = 0.19, and 95% confidence interval of [0.04, 0.12]. Tests showed no effect of iteration for either head or mantle regions: Mantel r = 0.002, two-tailed p = 0.97, and 95% confidence interval of [-0.03, 0.04] for the head region and Mantel r = −0.006, p = 0.92, and 95% confidence interval of [-0.11, 0.04] for the mantle region. A Mantel correlogram was computed to examine the underlying structure of the data where pixel series distance correlated significantly with animal identity. We found that REM-like state iterations that were more similar could be attributed to the same animal with Mantel r ~ +0.14 and that iterations that were more dissimilar could be attributed to different animals with Mantel r ~ −0.07 (Figure 5).

**Figure 3.**
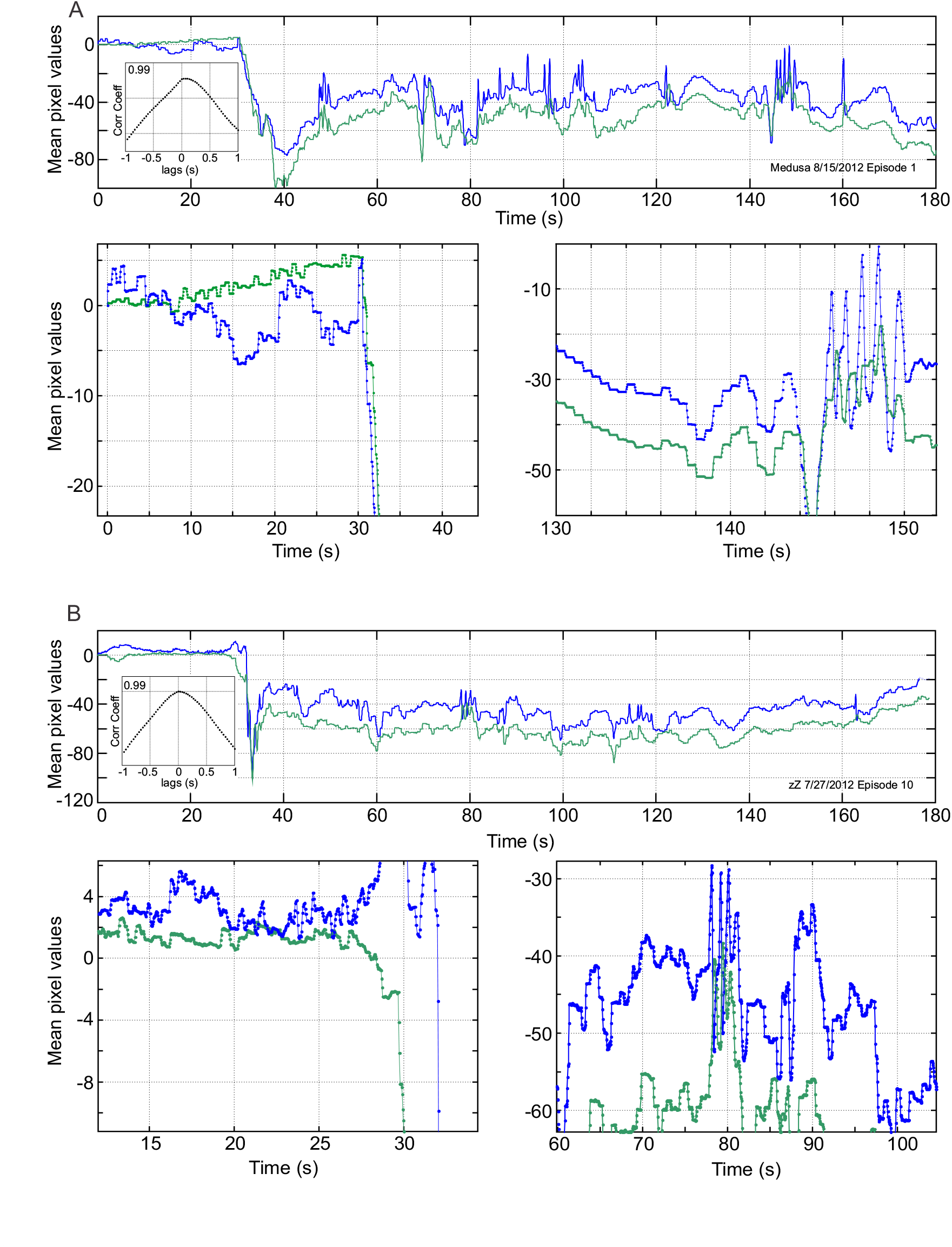
Two REM-like state iterations demonstrating common patterns of rapid brightness changes. Top panels in (A) and (B) show the full-length sequence for the head (green) and the mantle (blue) and bottom panels show expanded time series of the rapid onset of the REM-like state (left panels) and later in the REM-like state (right panels) as indicated by the x-axis time scale. Insets in top panels show the cross-correlation function between head and mantle time series. See Results for detailed explanation.

**Figure 4.**
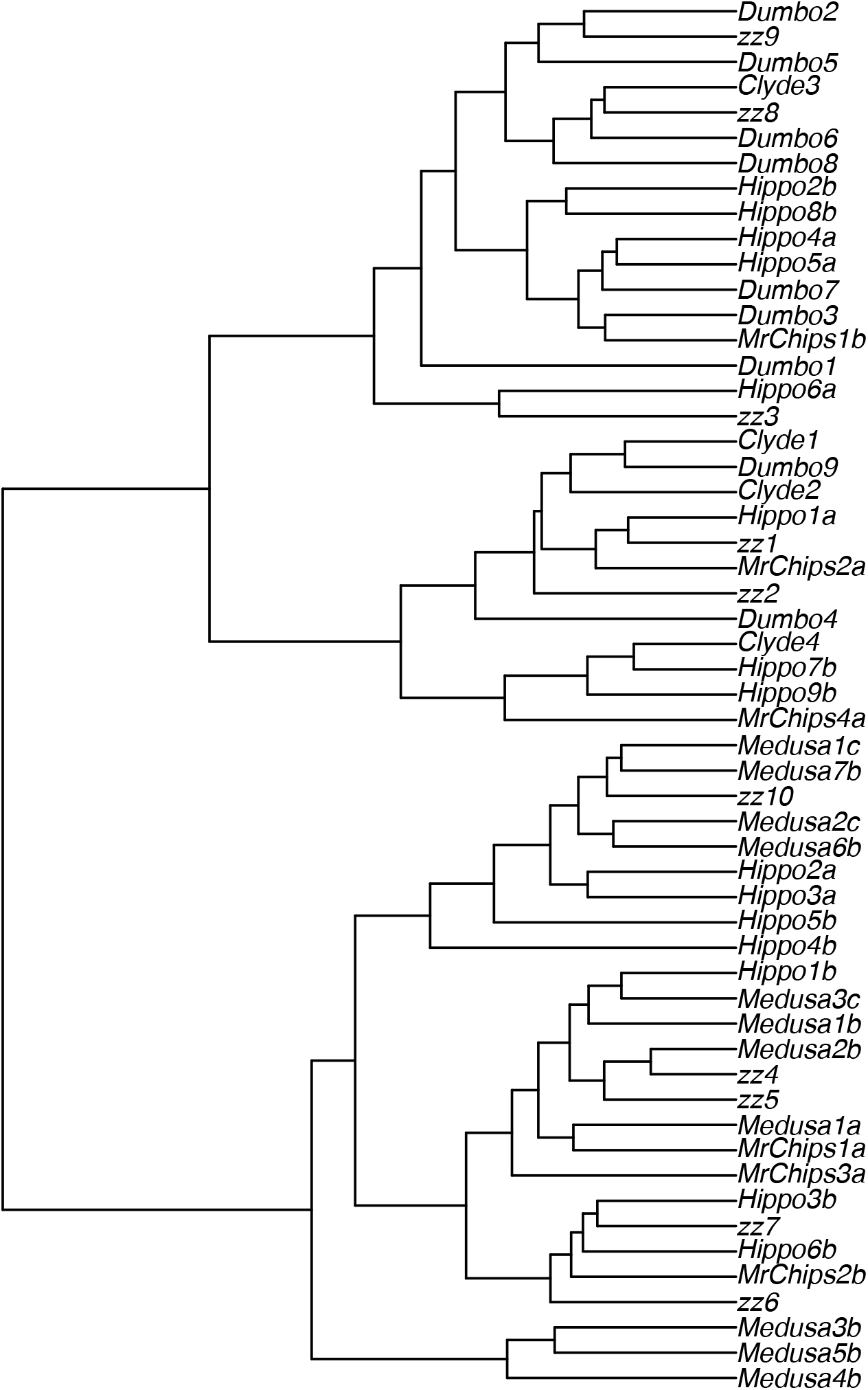
Cluster dendrogram of similarity between all REM-like state iterations using head data. Tip labels indicate the animal (H=Hippo, Z=zZ, P=MrChips, M=Medusa, D=Dumbo, and C=Clyde) and REM-like state iteration (1=1^st^, 2=2^nd^ … 10= 10^th^). Letters following the number indicate the sleep-like bout (a= 1^st^ bout, b=2^nd^ bout, c=3^rd^ bout).

**Figure 5.**
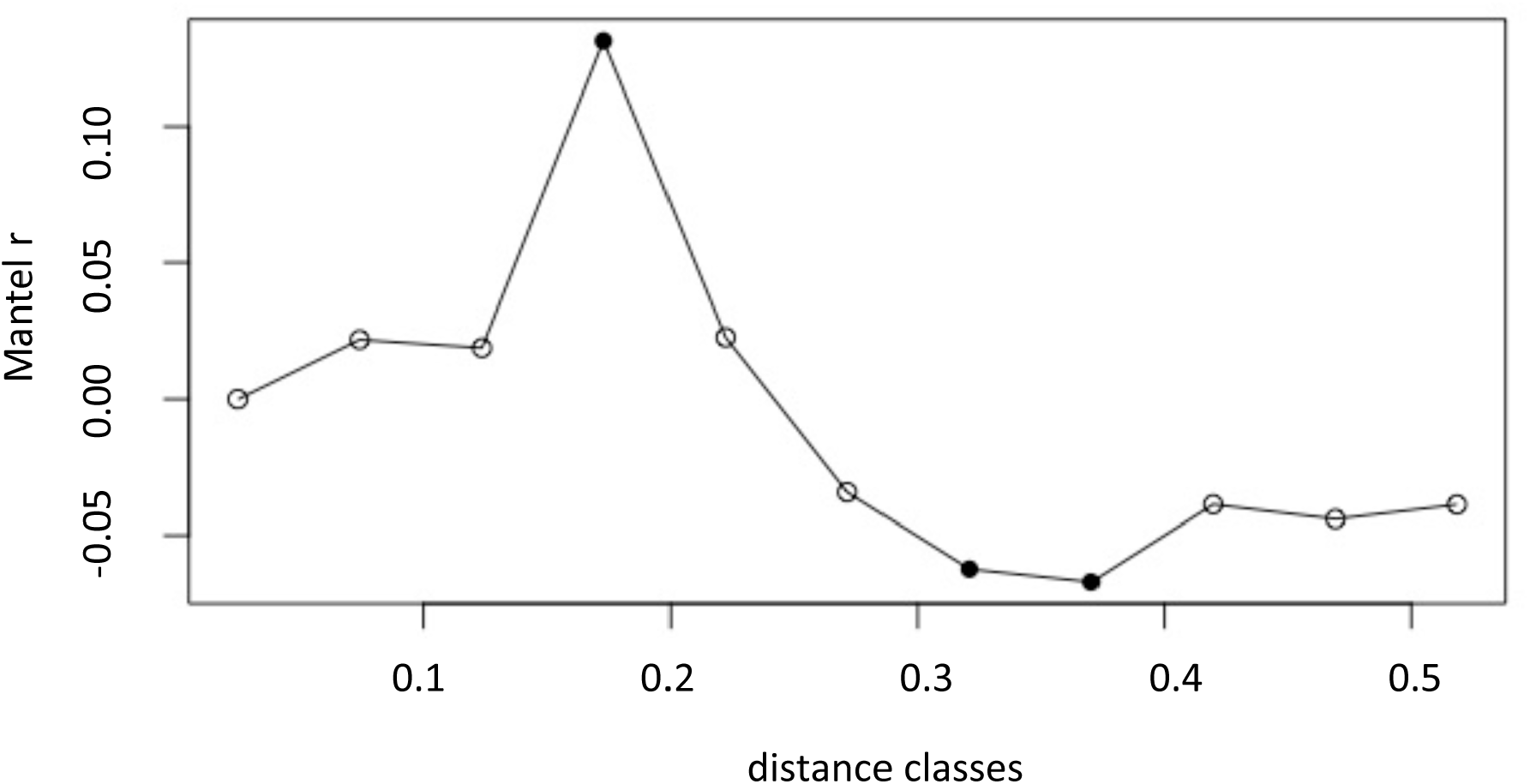
Mantel correlogram demonstrating the correlations between an individual animal and the pixel sequence distance matrix for all REM-like state iterations. Dark circles indicate p < 0.05 significance. The points of significance indicate that REM-like state sequences with higher similarity (larger Mantel r) originate from the same animal and sequences more dissimilar to each other originate from different animals. This relationship (Mantel r) is not very strong, however.

## Discussion

Slow-wave sleep and REM sleep states have been demonstrated in birds, mammals (Jerome M. Siegel, 2008) and reptiles (Shein-Idelson et al., 2016). Here we show that cuttlefish in a quiescent, sleep-like state periodically undergo a REM-like state, characterized by (i) general immobility with occasional muscle twitching (arms and neuromuscular chromatophore organs), (ii) rapid horizontal movements of the eyes, and (iii) alternating quiescent-sleep and REM-like states in a predictable ultradian rhythm. In octopuses, similar observations of quiescent behavior have been reported, including (i) circadian rhythms in activity, (ii) quiet periods characterized by closed eyes and a lack of activity, (iii) rebound when quiescence was prevented, and (iv) heightened sensory thresholds during quiet periods (Brown et al., 2006; Meisel et al., 2011). There are also anecdotal reports in octopuses of body patterning inconsistent with camouflage during this quiescent state (Meisel et al., 2011). If borne out by more quantitative measures (behavioral and neurobiological), collectively this would suggest that some cephalopods including the common cuttlefish, *Sepia officinalis*, and possibly the common octopus, *Octopus vulgaris*, experience a state akin to REM sleep.

Frank et al. (2012) described REM-like sleep behavior in adult, senescing cuttlefish but only observed a single iteration of the REM-like state per animal. In the present study, we recorded 55 iterations of REM-like behaviors in six non-senescing cuttlefish of the same species. The behaviors exhibited in both studies included spontaneous and rapid pattern and brightness changes, arm twitching, and eye movements. It is not implausible that the behaviors observed and reported in Frank et al. (2012) of senescing adults could have been a result of the deterioration of physiological systems that can affect sleep behaviors in senescent animals (Pandi-Perumal et al., 2002); especially since, as discussed below, such behavior was not detected in younger cuttlefish. However, our demonstration that the REM-like state occurs in adult, non-senescing cuttlefish nullifies this possibility.

Frank et al. (2012) also tested juvenile cuttlefish and observed homeostatic regulation of the QS state; however, they did not observe the REM-like state in juveniles. One possible explanation for this absence is neural immaturity. Alternatively, the REM-like state may be decreased or absent in situations that are stressful and demand increased vigilance. For example, most species show a circadian periodicity of REM; however, marine mammals, such as pinnipeds, show significantly decreased durations of REM sleep while in the water as opposed to when on land (O. Lyamin et al., 2008; O. I. Lyamin & Mukhametov, 1998) and migrating birds decrease REM sleep while in flight (Rattenborg, 2006; Rattenborg et al., 2016). Frank et al. (2012) also noted that the juvenile cuttlefish in their study may not have been provided a sufficiently long acclimation period. Technical issues prevented continuous recording of cuttlefish in our current study; however, the longest continuous recordings allowed us to observe two animals for more than 24 consecutive hours (Hippo: 53 hours; zZ: 40 hours). For the duration of these videos, the animals did not experience REM-like states but did appear to spend time in the QS state. Because the REM-like state was observed in Hippo prior to this time block, it suggests that the animal was acclimated adequately to allow this behavior to occur. If we allow that stress effects are not responsible for altering the expression of the REM-like state in these animals, an alternative explanation may be that *Sepia officinalis* may not experience this state every 24-hours as other animals experience true REM. Future behavioral study in juvenile and adult cuttlefish will be necessary to explore these possibilities.

Pixel intensity changes in small patches of chromatophores recorded in REM-like state iterations were not identical within individuals, nor were they identical between animals (Figure 4, Figure S3). Moreover, the expressions of individual chromatic components, as well as whole body patterns, were always highly transitory and often different from the way they are expressed in active cuttlefish. They were expressed in a seemingly disorganized manner; i.e. a typical camouflage pattern such as Mottle was shown often but briefly, whereas in active animals this pattern is maintained for many minutes or hours. The Mottle pattern was only shown acutely in cuttlefish in the REM-like state, however, and was preceded or followed by patterns that function as conspicuous displays such as Deimatic and Passing Cloud (Adamo et al., 2006). There were some similarities among REM-like state iterations (Video S6), however. For example, all animals performed the Passing Cloud display and most performed symmetrical/asymmetrical activation of mantle deimatic spots. These displays are associated with antipredator behavior and are therefore normally expressed in response to a perceived threat (Hanlon & Messenger, 1988; Langridge, 2006). None of the animals were in visual contact with other animals or with humans; it is possible — but highly unlikely—that the body pattern changes seen during the QS state in these experiments were a result of artefactual stimulation outside of the experimental chamber, such as ground-transmitted vibrations. Given that the system had a continuous inflow of local seawater, it is also possible that a chemical cue was responsible for eliciting these body patterns; however, all tanks received the same water and the REM-like state was not observed in multiple cuttlefish simultaneously. Rather, it is more likely that the cuttlefish were indeed exhibiting a typical behavior associated with the sleeplike state.

We recognize our methods may underestimate the similarity in gross pattern performance within and between animals because we compared pixel intensity changes for relatively small areas of the animal over temporally short periods of time. The animals actually performed many recognizable components of their patterning repertoire as mentioned above, but did not express these components in the same order or at the same time across REM-like state iterations. For example, all individuals displayed, either bilaterally or unilaterally, the 3^rd^ pair of deimatic spots on the mantle; however, the appearance of the spots did not occur at similar time points within the REM-like state iteration or even in all REM-like state iterations by the same individual (Video S6). The only recognizable common chromatophore pattern that was displayed in a temporally predictable manner was the presence of the white head bar, which turned dark and was accompanied by the eye squint, marking the start of the REM-like state. All other components were recognizable but the order in which they were performed and the duration of appearance was variable. Many of these components are normally expressed in response to predators or perceived threats or are used in camouflage, yet the testing environment was static with regard to substrate, structures and visual cues.

Our findings suggest that during the REM-like state, fragments of normal waking activity in these areas are spontaneously re-activated, but in combinations not usually observed in wakefulness. This is reminiscent of mammalian REM sleep, which also contains fragments of waking patterns of activity in motor, limbic and cortical circuits. This is revealed most dramatically in mammals that lack the normal paralysis of REM sleep. Under such conditions, during REM sleep animals will display components of complete waking behaviors, including aggressive displays, orientation and stalking behavior (Dumoulin Bridi et al., 2015; Louie & Wilson, 2001; Morrison, 1979). Moreover, in REM sleep remote and recent memories are accessed and combined in complex ways not typically experienced in the waking state (Hobson, Pace-Schott, & Stickgold, 2000). While speculative, it is possible that the unusual combinations of chromatophore activity during REM-like states reflect a similar process in *Sepia officinalis*.

Although not the prime target of this study, the dynamics of brightness changes would be worth investigating more quantitatively. Of particular note, (1) all REM-like state iterations begin with a rapid darkening that is synchronized between head and mantle regions of interest (bottom left panel in (Fig. 3A), but can also involve delays of several seconds (bottom left panel in (Fig. 3B)); (2) overall, brightness changes in the head and the mantle region are correlated, often with a delay of 30 to 100 ms (insets in top panels in (Fig. 3A and B); (3) such correlations are not present, however, in the fine-scale brightness change on short time scales (Fig. 3, bottom panels); (4) rapid brightness changes occur in discrete steps, even before the REM-like state starts (Fig. 3, bottom panels), but (5) can also involve bursts of smooth oscillatory changes (Fig. 3, bottom right panels); (6) both head and mantle regions tend to brighten slowly throughout the course of a REM-like state iteration (top panels Fig. 3A and B). Given that these brightness changes reflect chromatophore dynamics and the activity of neural networks, it would be interesting to compare the temporal organization of brightness changes in awake and “sleeping” cuttlefish and to eventually relate them to changes in chromatophores and to the neural activity causing them.

If future work reveals cephalopod QS state and the REM-like state to be analogous to vertebrate slow-wave and REM sleep, then it is worth remarking here that the periodicity of the REM-like state in the cuttlefish appears more similar to the periodicity of REM sleep in small mammals than to birds or reptiles. The total average time an animal spent in the REM-like state across a single QS-REM-like state cycle is similar to small mammals and birds. Regardless of whether sleep in animals resulted from single or multiple evolutionary origins, this phylogenetic distribution of patterns of periodicity, total duration of specific sleep-states, and flexibility to delay REM sleep (or REM-like states), promises a fruitful avenue towards identifying ecological pressures that shape sleep behavior (Aulsebrook et al., 2016).

The possible occurrence of sleep and REM-like sleep in a clade of organisms that is distantly related to any clade where REM sleep has been observed previously poses a challenging and important opportunity to broaden our understanding of sleep and REM sleep (Corner, 2013; J. M. Siegel, 1995; Irene Tobler, 1988). Cephalopods have the largest and most complex brains — and the most diverse repertoire of behaviors — of any invertebrate animal (Hanlon & Messenger, 2017; Nixon & Young, 2003). Whether future work demonstrates that cuttlefish adhere to all the criteria deemed necessary for a legitimate sleep state, or prove to be an exception to the general pattern described in the literature for vertebrates, examining these questions in an organism that is ecologically distinct and phylogenetically distant from current model organisms should serve to inform the broader questions of how to accurately define sleep, discover its functions, and understand its evolutionary origins.

## Acknowledgments

We thank the Grass Foundation for support and the summer 2012 MBL Grass Laboratory Directors and Fellows for early feedback and assistance. We thank the staff at the MBL for their indispensable technical and logistical assistance. Thanks to members of the Hanlon lab for culturing the cuttlefish. JGB thanks the Millersville University Faculty Grants Committee and RTH thanks the Sholley Foundation for partial support.

